# Parameter-dependent shift from rational to irrational decisions in mice

**DOI:** 10.1101/701722

**Authors:** Nathan A. Schneider, Benjamin Ballintyn, Donald Katz, John Lisman, Hyun-Jae Pi

**Author notes:** Correspondence: Hyun-Jae Pi. **Author Contributions:** H.P. initiated and oversaw the project; N.S., D.K., J.L. & H.P. contributed to designing the behavioral task; N.S. performed experiments; N.S., B.B, & H.P. analyzed data; B.B. performed a theoretical analysis and computational modeling; N.S., B.B., D.K. & H.P wrote the manuscript.

## Abstract

In the classical view of economic choices, subjects make rational decisions evaluating the costs and benefits of options in order to maximize their overall income. Nonetheless, subjects often fail to reach optimal outcomes. The overt value of an option drives the direction of decisions, but covert factors such as emotion and sunk cost are thought to drive the observed deviations from optimality. Many questions remain to be answered as to 1) which contexts contribute the most to deviation from an optimal solution; and 2) the extent of these effects. In order to tackle these questions, we devised a decision-making task for mice, in which cost and benefit parameters could be independently and flexibly adjusted and for which a tractable optimal solution was known. Comparing mouse behavior with this optimal solution across parameter settings revealed that the factor most strongly contributing to suboptimality was the cost parameter. The quantification of sunk cost, a covert factor implicated in our task design, revealed it as another contributor to suboptimality. In one condition where the large reward option was particularly unattractive and the small reward cost was low, the sunk cost effect and the cost-led suboptimality almost vanished. In this regime and this regime only, mice could be viewed as close to rational. Taken together, our findings support a model whereby parallel neural circuits independently activate and modulate multiple valuation algorithms, and suggest that “rationality” is a task-specific construct even in mice.

**Significant Statement:** Irrational factors in economic decision-making often cause significant deviation from optimal outcomes. By devising a flexible economic choice behavior for mice and comparing their behavior with an optimal solution, we investigated overt and covert factors that contributed to suboptimal outcomes and quantified the deviation from optimality. This investigation identified regimes where mice could be viewed as rational or irrational depending upon the parameters in the same task. These findings may provide a platform to investigate biological substrates underlying rational and irrational decision factors.

## Introduction

The classical models of economic decision-making, particularly as applied to foraging behavior, present subjects as rational agents who maximize gain by deliberately analyzing the cost and benefit associated with available options (1–6). The principle of this optimization has provided a theoretical foundation capable of explaining and predicting a plethora of phenomena in decision-making.

Often, however, the outcome of the decision-making deviates significantly from the optimal (7–10). This is because the value assessment of given options is continually influenced by the subject’s fluctuating intrinsic state and a constantly changing external environment (11–13). While the overt value of the given options, such as the quantity or the calorie content of food, are inevitably major forces driving the direction of the choices, models incorporate covert factors to explain the frequent deviations from optimality (7, 14, 15). Much progress has been made in understanding these factors, but questions still remain as to which contexts most drive the deviations from optimality, as well as to the context-dependence of the deviations’ magnitude.

Although research using non-human laboratory animals has provided vast opportunities for investigating the biological substrates and neural mechanisms underlying decision-making, the non-verbal nature of these subjects makes topics such as suboptimality and irrationality difficult to study. Nonetheless, a series of recent studies by the Redish group applied quantitative behavioral readouts and theoretical frameworks, to investigate covert decision factors in laboratory animals (9, 14, 16, 17). These studies reveal the influence of the sunk cost fallacy, an irrational thought process wherein a subject tends to continue a behavior based on previous investment choices even if the decision is unprofitable overall (18, 19). One study identified a similar behavioral signature of sunk cost in humans, rats, and mice (9).

We sought to further understand the impact of the overt and covert factors on decision-making by extending this approach in a mouse model system. Bringing the above-described work together with several past studies on decision-making in the context of foraging behavior (17, 20–26), we devised an economic choice task in which cost and benefit parameters can be flexibly and independently adjusted, and in which a tractable optimal solution is always identifiable.

In our behavioral task, on each trial mice make the decision to lever press at a progressive or fixed ratio reward option (PR; FR) with different amounts of water reward associated with each (21); the shift of ratio preference was quantified by indifference points where the values of the two sides became equivalent. We tested whether mice adjusted their decisions and indifference point depending on the cost-benefit parameters, and found that their switching decisions did indeed shift accordingly. By comparing the actual behavioral outcomes with the optimal solutions, a deviation from optimality could be quantified and compared across parameter settings.

Doing so, we found that differences in the cost parameter had a more significant impact on suboptimality than the benefit parameter. Mice exhibited behavior similar to that observed in sunk cost studies (9, 19), and indeed, further analysis revealed that sunk cost contributed to the suboptimal outcomes. This contribution was highly parameter-specific, however, in that we identified a regime in the parameter space where the cost parameter and sunk cost had no significant impact. Therefore, mice can be viewed as rational agents in some regimes, and not in others – a human-like behavior. Thus, our approach provides a platform to investigate both rational and irrational decisions in the same task.

## Results

### A flexible economic choice behavior for mice

In order to quantitatively evaluate the factors that influence economic decision-making, we devised a behavioral paradigm that allowed us to flexibly and independently adjust the costs and benefits of the possible choices. In addition, this task has a tractable optimal solution that provided a comparison of how close mouse behavior was to optimal.

Water-restricted mice were required to press levers to collect a water reward (Fig. 1A). Mice could freely choose to press either a fixed ratio (FR) lever or progressive ratio (PR) lever. The FR lever provided a small volume of water reward each time mice completed a fixed (i.e., unchanging) number of presses (Movie S1); the PR lever, meanwhile, provided a large volume of water, but achieving this volume required a number of presses that increased (e.g., 2, 3, 4…) with each successful reward (Movie S2). Mice pre-trained to press both levers reliably preferred the PR lever at the start of each session, undoubtedly because this strategy afforded them a large reward with little effort compared to the FR (Fig. 1B). As the session progressed and the PR requirement became higher, however, this cost (i.e., the required number of presses) overwhelmed the benefit of the large reward, and mice switched their preference to the FR lever, supporting the general hypothesis that mice evaluate the relative values of given choices in terms of both costs and benefits.

**Fig. 1.**
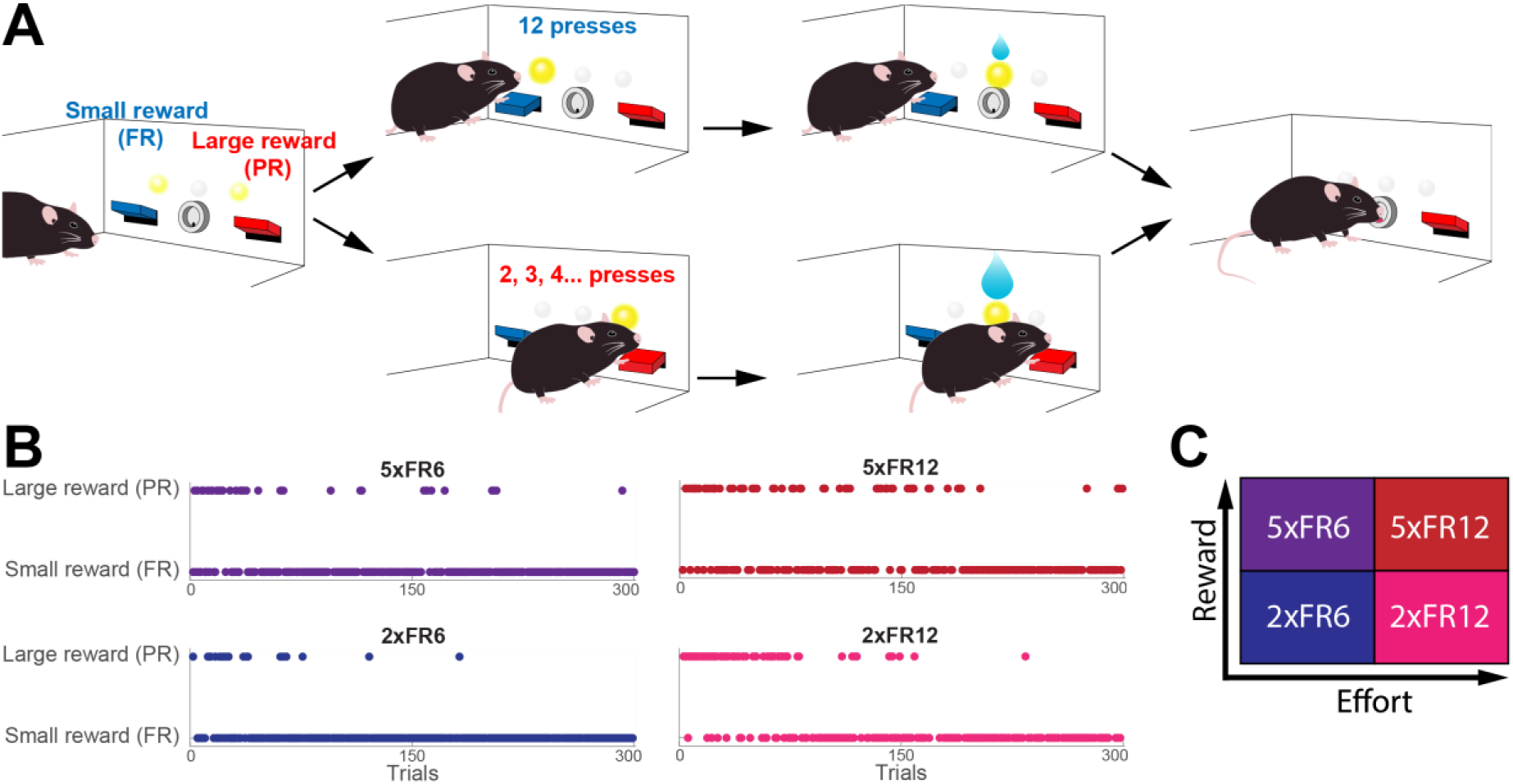
Optimal Switching Behavior Task. **A**. Schematic of task design. **B**. Example sessions for each parameter pair. Each point is a single trial within the session. Only the first 300 trials of each session are shown. All sessions are from the same animal. **C**. Grid showing increasing reward and cost for each type of parameter pair.

### Mice adjust switching decisions proportional to the values of session parameters

Having established the basic rationality of task performance, we next asked how the changes in relative reward size (the benefit parameter) and the number of lever presses (the cost parameter) affected the choice behavior. By fixing one parameter (e.g., reward amount) and changing the other (e.g., lever press requirement), the contribution of cost and benefit in decision-making can be evaluated semi-independently within the same task, across a wide range of parameter combinations. In this initial study, we chose four combinations of parameters by crossing the ratio of large reward to small reward (2:1 or 5:1) with the number of presses required for the FR reward (6 or 12) and devised a nomenclature to differentiate the parameters. For instance, 2xFR12 means the volume of large reward, 6 μL, is twice as much as that of the small reward, 3 μL, and the FR requirement is 12 presses; the four combinations of parameters can be denoted as 2xFR6, 2xFR12, 5xFR6, and 5xFR12 (Fig. 1C).

We tested the hypothesis that changes in the relative values of given choices would yield quantifiable differences in the switching decisions of 10 mice exposed to each of these parameter combinations. An equal number of male and female mice were evaluated, but as no significant male-female differences were not observed (Fig. S1A); we combined the data for the below analyses. Four sessions with each parameter combination were collected (160 sessions total: 4 sessions × 4 parameter pairs × 10 mice). Left and right levers were pseudo-randomly assigned to be PR and FR at the start of each session.

Consistent with our initial observation, mice initially preferred the large-reward PR under all parameter settings and switched their preference to the small-reward FR as the number of presses required for the PR reward increased (Fig. 1B). All mice were able to make switching decisions by accurately evaluating the cost-benefit relationship of the given choices.

The point at which switching decisions were made varied between conditions, however, a fact that is visible in a simple plot of decisions over time (Fig. 1B). The switch to FR happens earliest in the 2xFR6 condition and latest in the 5xFR12 condition, assuredly because of the different relative values of the PR. Under the simple assumption that these parameter variables directly manipulate the relative value of the PR, the value of the large-reward PR is lowest at 2xFR6 because the PR reward is low and the effort and time costs for the FR are also low. Alternatively, the PR value is highest at 5xFR12 because the reward is high and the alternative choice requires more effort and time. The PR values of 2xFR12 and 5xFR6 are somewhere in between the other two. Reflecting on these relative values, the number of PR choices increased as the value increased. These measures were quantified in percentage and deviation of the PR choices (Fig. 2A, B). The percentages of choosing the PR with large reward were 6.5 ± 1.4%, 14 ± 3%, 7.8 ± 1.5%, and 19 ± 4% (mean ± SEM) at 2xFR6, 2xFR12, 5xFR6, and 5xFR12, respectively. The percentage of the FR choices were 86%, 69%, 84%, and 63%, respectively (Fig. 2A).

**Fig. 2.**
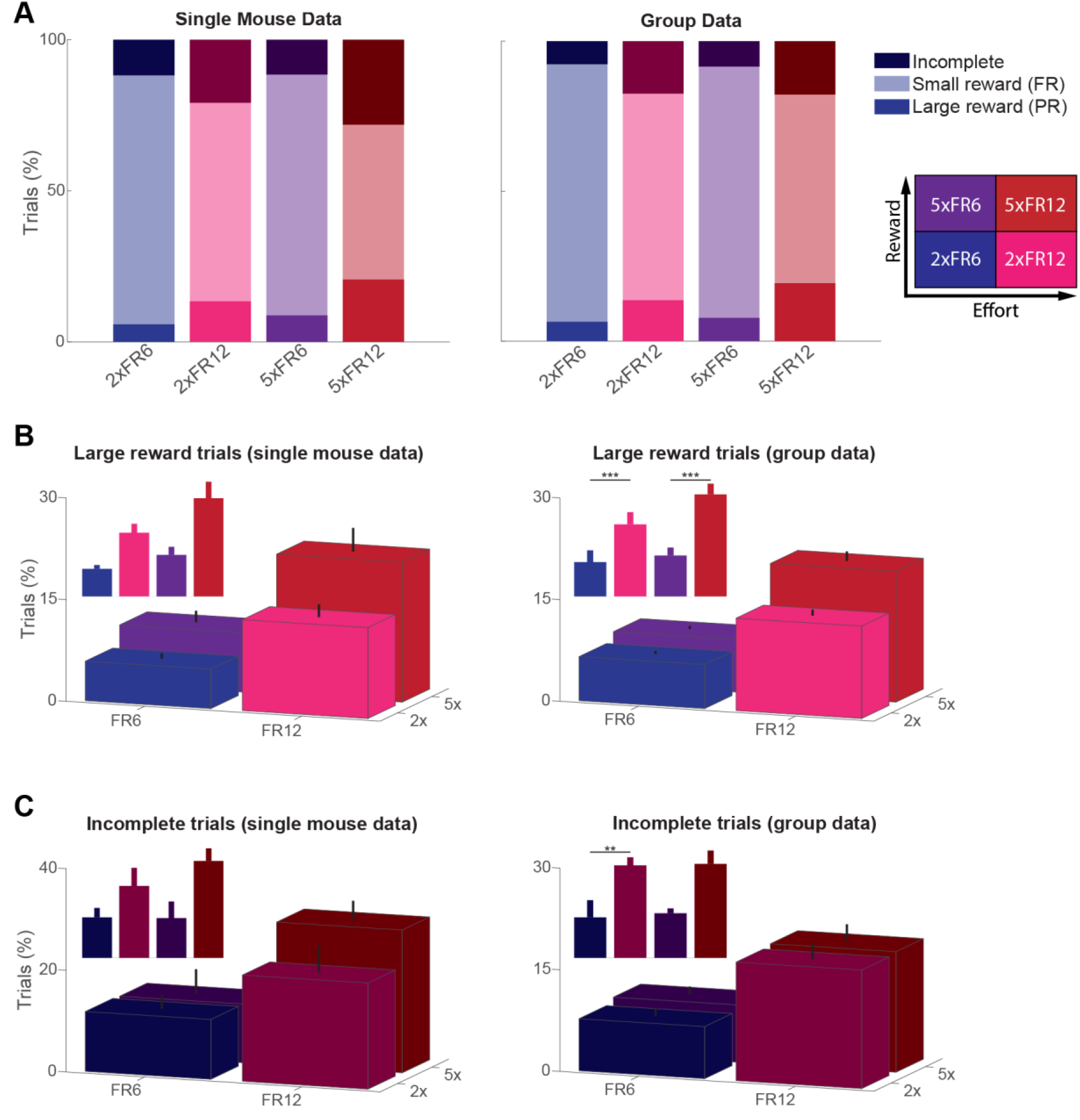
Trial type ratios reflect evaluation of changing cost and benefit between session parameters. **A**. Stacked bar graph showing trial distributions for each parameter (left, n = 4 sessions; right, n = 10 mice). **B**. Percent of trials where large reward (PR) was collected (left, n = 4 sessions; right, n = 10 mice). Error bars reflect the standard error of the mean. A two-way Scheirer-Ray-Hare test indicated a significant effect in the FR requirement (H_1_ = 28.98, p = 7 × 10^−8^). Insert shows the same data with significant pairs notated (post hoc Wilcoxon Rank-sum test with Bonferroni correction, \*\*\**P* < 0.001). **C**. Percent of trials where reward collection was failed (left, n = 4 sessions; right, n = 10 mice). Error bars reflect the standard error of the mean. A two-way Scheirer-Ray-Hare test indicated a significant effect in the FR requirement (H_1_ = 16.46, p = 5 × 10^−5^). Insert shows the same data with significant pairs notated (post hoc Wilcoxon Rank-sum test with Bonferroni correction, \*\**P* < 0.01).

An analysis of the percentages of trials that mice failed to complete provided data consistent with the above conclusions (a trial was considered as incomplete if the mouse paused activity for more than ten seconds, or commenced in pressing the alternative lever, after initiating a trial but before completing the total number required. If the relative values of the PR side were higher, mice tried and failed more (Fig. 2C) and the failure was more frequent on the PR side (Fig. S1C, S1D). The incomplete trial percentages were 8 ± 3%, 17 ± 7%, 8 ± 4%, and 18 ± 8% for 2xFR6, 2xFR12, 5xFR6, and 5xFR12 respectively. Increasing the cost parameter (i.e. number of FR presses) contributed more to the incomplete trials. The parameters 2xFR6 and 5xFR6 have a difference of only 0.7%, yet between 2xFR6 and 2xFR12 there is a difference of 9% (p < 0.01, Wilcoxon Rank-sum test with Bonferroni correction). The cost change also exerted a stronger effect on the task performance as indicated in the total number of trials and water collected (Table S1). Consistent with our hypothesis, these results suggest that mice can differentiate the relative values of the session parameters and adjust their switching decisions accordingly.

### Quantification of switching decisions by indifference points

One idea implemented in our task design was utilizing an “indifference point” as a behavioral readout of how mice evaluate cost and benefit. Theoretically, these points occur when the subjective values of each side become equal. In our behavior task, initially, the PR with large reward is more valuable than the FR with small reward. As the PR requirement increases, however, the value on the PR side starts to decrease while the FR value remains fixed. At some point, the subjective value of the PR becomes equivalent to that of the FR:

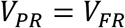

The PR requirement at the indifference point can also be interpreted as mice’s willingness to work to collect large reward. We estimated the PR requirement at indifference points by fitting the experimental data with a function that represented the distribution of the choices. Because our data was binary (i.e. two choices), session data were fit with a sigmoid function (Boltzmann function). Sigmoid fitting curves captured the profiles of mouse decisions, showing the transition from the PR to the FR (Fig. 3A, B). The indifference trial number was first estimated where the sigmoid curve crossed the midline. Then the number of lever presses required for the PR at that trial was extracted from the data, which provided the PR requirement at the indifference point of the session data. Fig. 3A also shows where the indifference point lies compared to the increasing PR requirement over time. The results showed that both at a single mouse level and at the animal average, the estimated PR requirement at indifference points was lowest at 2xFR6, highest at 5xFR12 and those of 2xFR12 and 5xFR6 lied in between (20 ± 4 for 2xFR6, 28 ± 4 for 2xFR12, 29 ± 4 for 5xFR6, and 45 ± 7 for 5xFR12; Fig. 3C). We also estimated indifference points using a median of trials which captured the spread of choices better and the results are presented in Fig. S2. Despite differences in the numbers, the two approaches showed the consistent shifting patterns of the PR requirement at the indifference point proportional to the relative values of the PR. This indicates that mice were able to adjust their switching decisions according to the given value.

**Fig. 3.**
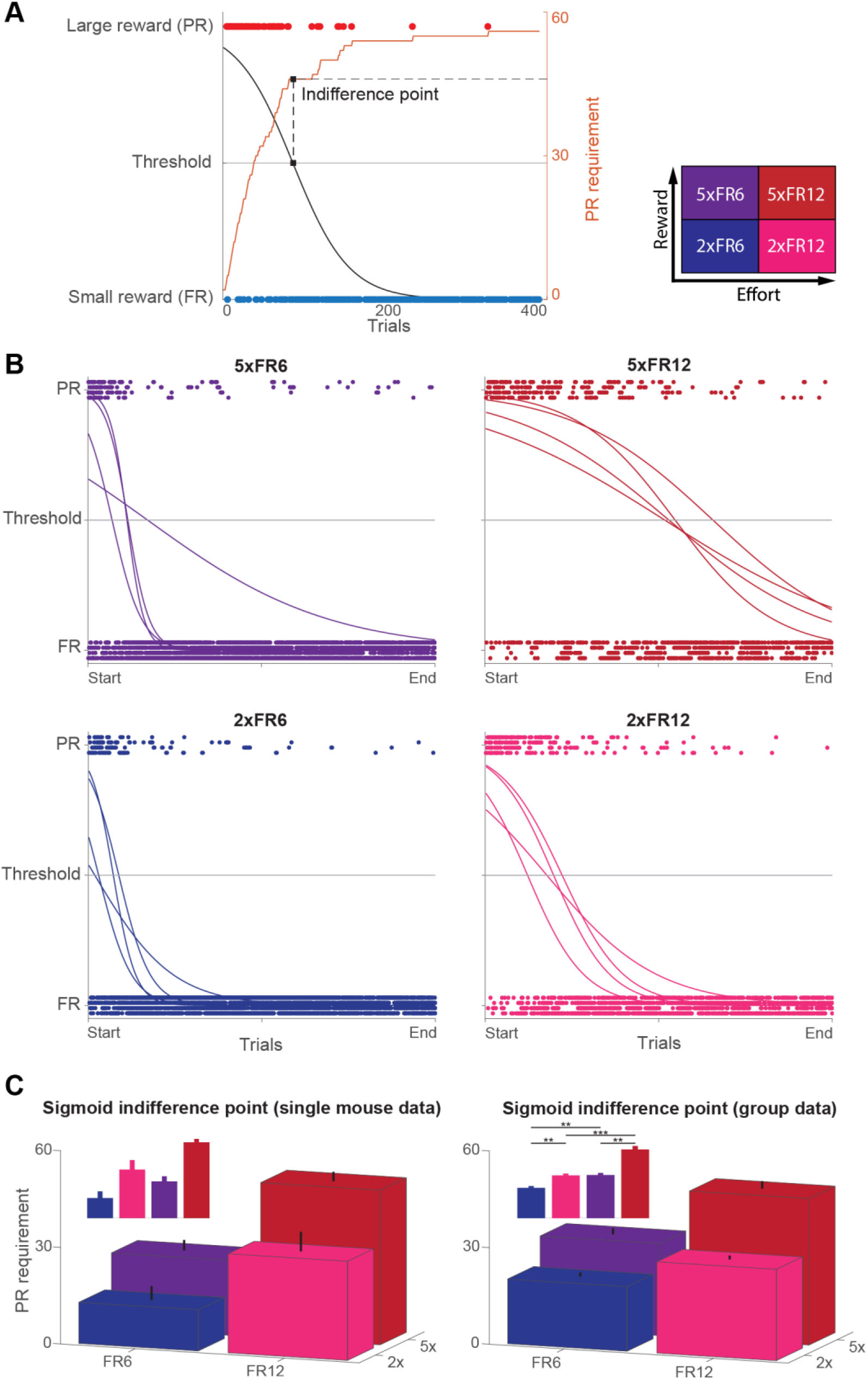
Estimating indifference points through a sigmoid curve shows variation in switching behavior between parameter conditions. **A**. Example session showing how the indifference point is estimated. Black curve is the sigmoid fit and orange curve is the cumulative PR requirement over time. First, the data is fitted to a sigmoid curve. Then, where the curve crosses the threshold, a trial number is identified (black squares). Finally, the indifference point is determined by finding the number of presses required for the PR at this trial (dashed lines). **B**. Sigmoid curves for all 16 sessions of one mouse, sorted by session parameters. Trials are plotted as individual points normalized to the total number of trials in the session and each row of points is a different session. For sessions with higher reward or effort, the threshold intersection shifts farther to the right. **C**. Indifference points for a single mouse (left, n = 4 sessions) and the entire population (right, n = 10 mice). Error bars reflect standard error of the mean. A two-way Scheirer-Ray-Hare test indicated a significant effect in the FR requirement (H_1_ = 14.05, p = 0.0002) and the PR reward volume (H_1_ = 15.72, p = 7 × 10^−5^). Insert shows the same data with significant pairs notated (post hoc Wilcoxon Rank-sum test with Bonferroni correction, **p < 0.01, ***p < 0.001).

### Cost parameter contributes more significantly to the suboptimality

When is the right moment for mice to switch their preference from the PR to the FR? What is an optimal strategy to maximize gains? How close to the optimal is mouse behavior? In order to answer these questions, we took advantage of optimality models in foraging theory. The long-term rate of gain intake (also referred to as the ratio of expectations or RoE) is often assumed to be the optimal ‘currency’ to maximize because it minimizes the loss of alternative opportunities (27). An alternative to RoE, which is not theoretically optimal but may better represent the currency that animals actually maximize, is the expectation of ratios (EoR, also called the short-term rate) which measures the average per-trial ratio of gains to costs (27). The comparison between RoE and EoR analysis in our task is described in “Maximizing of different currencies yields the same optimal behavior” in SI Appendix.

Because of the trial-based structure of the task, we studied mouse behavior in reference to the EoR model where a rational agent is assumed to maximize the average per-trial rate of benefit over cost. EoR is given by the following equation:

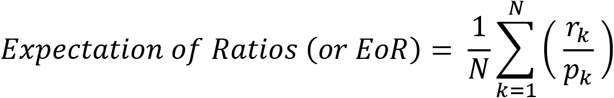
where *N*, *r*_*k*_, and *p*_*k*_ denote the total number of trials, reward on trial *k*, and cost of lever-press on trial *k*, respectively.

Using this equation, we first calculated the optimal trial numbers that the mouse should spend at each side for a given session type. Given a certain number of trials, we asked what was the optimal distribution of the choices between the PR and FR side where optimality means maximizing the EoR. The estimated optimal numbers of trials at the PR side 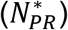 are 10, 22, 28, and 58 trials at 2xFR6, 2xFR12, 5xFR6, and 5xFR12 respectively (Fig. 4A; see EoR analysis in SI Appendix). Using these numbers, the optimal EoR (EoR^opt^) was calculated and compared with EoR^mice^. The EOR optimality was defined as the ratio of EoR^mice^ over EoR^opt^.

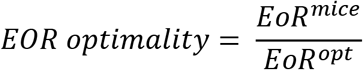

**Fig. 4.**
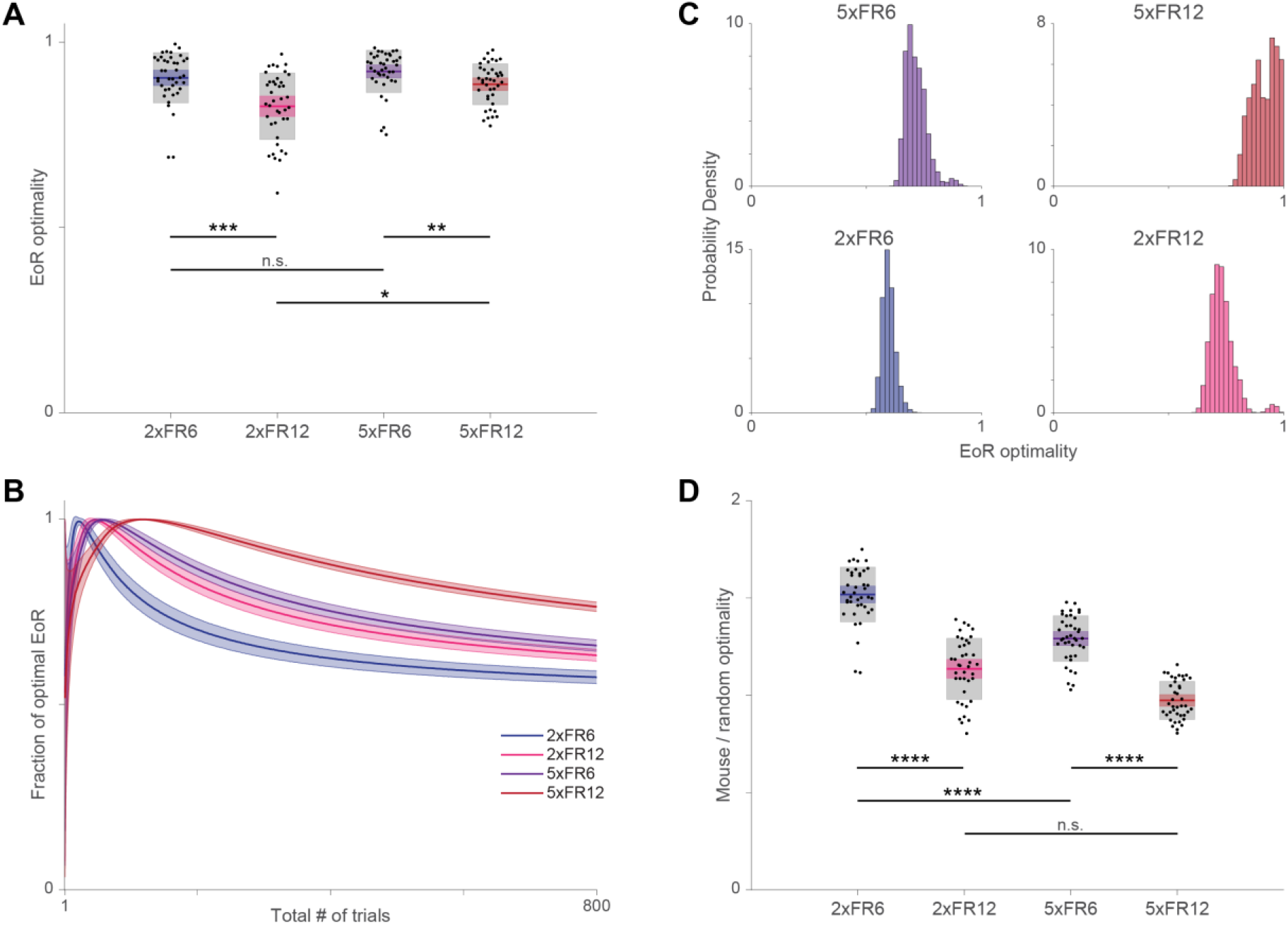
Rate-maximizing (EoR) and random choice models. **A.** EoR optimality distributions by session type. Each point represents the EoR optimality from one session. Colored line, colored shaded box, and grey box show the mean, 95% confidence interval for the mean, and the standard deviation, respectively. Scheirer-Ray-Hare test indicates a significant effect of both PR reward size and FR lever press requirement on EoR optimality .*p < .05, **p < .01, ***p < .001 indicates significance for post-hoc pairwise two-tailed Wilcoxon rank sum tests. **B.** EoR optimality of a randomly choosing agent for a given number of trials. For a given number of trials (1–800) 10,000 agents that chose randomly between the FR and PR sides were simulated and their fraction of the optimal EoR for that number of trials was recorded. Dark lines indicate the mean and shaded regions show the standard deviation. **C.** Distribution of EoR optimality for randomly choosing agents given mouse trial statistics. For each session a mouse performed, 10,000 randomly choosing agents were simulated for the number of trials the mouse performed that session. Therefore, each histogram shows the results from 400,000 random agents. Because the domain of each histogram is smaller than 1, probability density of each bin can be greater than 1. **D.** Comparison of observed mouse EoR optimality to that of the random agent. For each session a mouse performed, it’s observed EoR optimality (shown in **A**) was divided by the mean EoR optimality of the 10,000 agents simulated for that number of trials (mean value for appropriate session type and number of trials shown in **C**). Scheirer-Ray-Hare test again showed significant effect of PR reward size and FR lever press requirement on this ratio. Significance bars for post-hoc tests are as in **A**. ****p < 0.0001 indicates for post-hoc pairwise two-tailed Wilcoxon rank sum tests.

This value lies between 0 to 1, where 1 means mice performs the task as an optimal agent.

The mean EoR optimality was 0.90 ± 0.01, 0.83 ± 0.01, 0.92 ± 0.01 and 0.88 ± 0.01 (mean ± SEM) at 2xFR6, 2xFR12, 5xFR6, and 5xFR12 respectively (Fig. 4A). Note that these numbers might be around the upper limit of the true values considering the limitations of the EoR model (see Discussion). One noticeable trend was that the change in the cost parameter had a more significant effect on the deviation from the optimality. For instance, given a fixed benefit parameter (2x or 5x), the EoR optimality of FR6 was higher than that of FR12 (Fig. 4A). Given a fixed cost parameter, however, the change in the benefit had less influence on the deviation (Fig. 4A). One explanation of this result is that increasing the cost parameter contributed more to the incomplete trials (Fig. 2A) that contributed negatively to the rate of reward collection. In summary, the cost parameter, not the benefit, was one main source that led to the suboptimal outcome.

### Sunk costs contribute to the suboptimality

The sunk cost fallacy describes an irrational decision in which a subject continues investing in a bad option despite the diminished overall return (18, 19). Although the sunk cost has been a difficult topic to study in animal model systems, a recent study with an elegant design overcame the challenge and identified a behavioral signature of sunk cost in humans, rats, and mice (9). In our task, after passing the indifference point, mice often revisited the PR side which required an unreasonably high number of presses and continued investing their lever presses to collect the large reward. This observation was similar to what Sweis et al. reported (9) and led us to investigate whether the signature of the sunk cost could be identified in our dataset and the effect could be quantified.

Data were analyzed in a similar way as described in the report with the number of PR presses as a variable. After the initial choice of the PR, the probability of collecting the large reward given how many presses remained was calculated and fitted linearly (Fig 5A). This was repeated for several different investment groups to show how the probabilities of collecting reward changed as mice continued pressing the PR lever. If there was a sunk cost effect in our behavioral task, we would expect that as mice invested more presses into collecting the large reward, the probability of collecting reward would increase, reflected by a decrease in the regression slope (Fig. 5B).

**Fig. 5.**
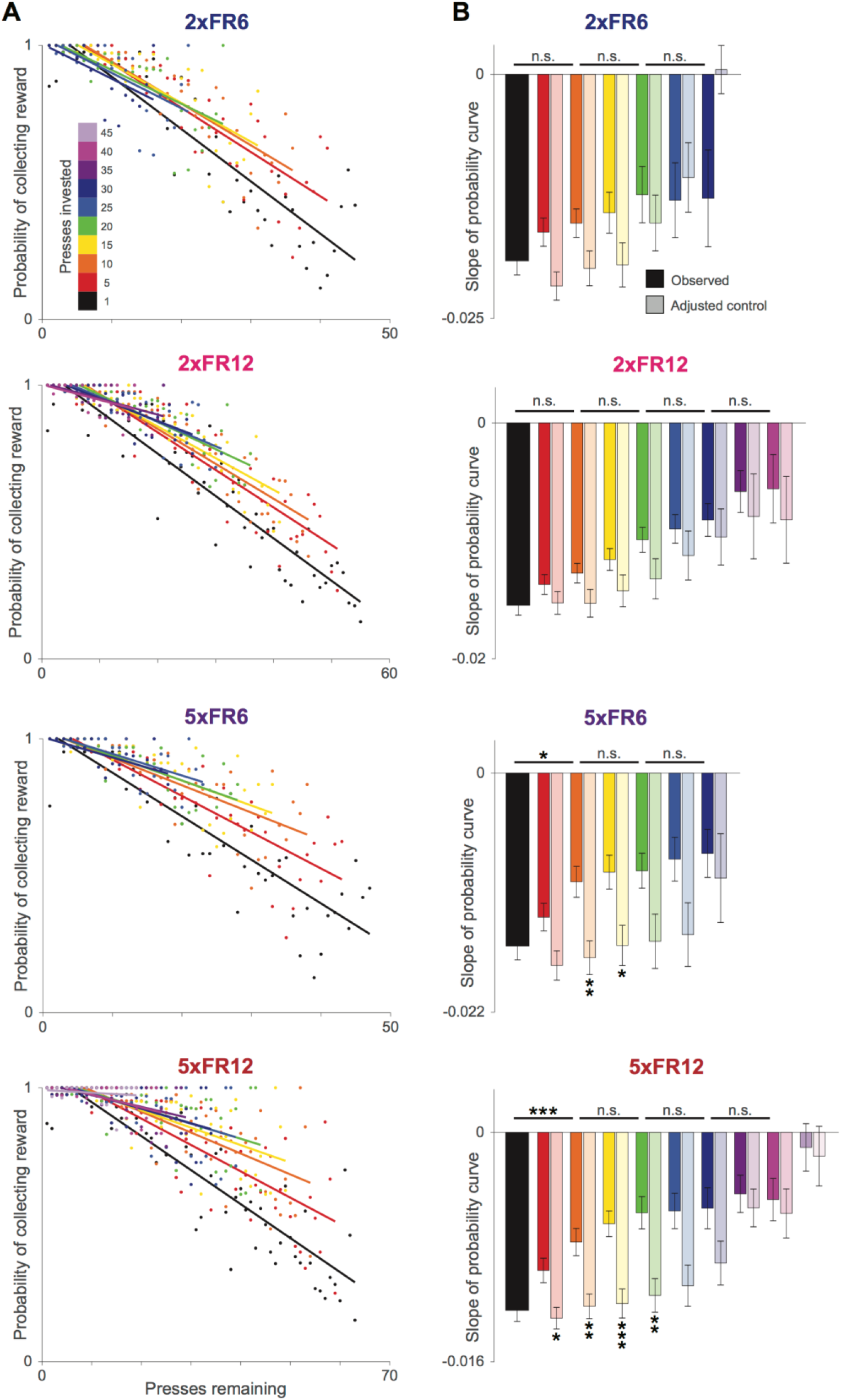
Sunk cost effect contributes to the suboptimality. **A.** Probability of collecting the large reward given how many presses are remaining and how many presses have already been invested. Each color reflects a different number of presses the mouse has already pressed. **B.** Slopes from each regression in **A** (“observed”) and slopes computed from black data points limited to the range of each colored group (“adjusted control”; Fig. S3). Error bars reflect standard error of the mean. An analysis of covariance (ANOCOVA) was used to compare slopes of the linear regression models, correcting for multiple comparisons. Not significant (n.s.), p < 0.05; *p < 0.05; **p < 0.01; ***p < 0.001; ****p < 0.0001.

The application of this analysis to our dataset revealed that indeed an effect of sunk cost existed. Surprisingly, it is not present in all four conditions of the task. A two-way ANOVA collapsing across sunk cost conditions revealed that the effect was only significant in contexts with a higher cost or benefit parameter (2xFR12: F = 31.8, p < 0.0001; 5xFR6: F = 8.4, p < 0.01; 5xFR12: F = 6.76, p < 0.01) and not in the lowest PR value context (2xFR6: F = 3.1, p = 0.08). The slopes of sunk cost conditions were also compared to neighbors and adjusted controls to illustrate that sunk cost effects increased with an increase in investment (e.g. 5xFR12 comparison of 1 press and 10 presses invested: p < 0.001). Taken together, the results suggest that the sunk cost fallacy also contributes to suboptimality.

### Optimal and suboptimal regimes in the parameter space

The results thus far indicated that whereas the cost parameter contributed to the suboptimality both overtly and covertly, the benefit parameter did so indirectly via sunk cost. Therefore, different cost and benefit parameters generated a different effect on each condition. Interestingly, however, neither sunk cost nor the cost-led suboptimality had much impact on 2xFR6 where the large reward option was particularly unattractive and the small reward cost was low. Taken together, in this condition, the performance of mice in the 2xFR6 condition might be close to that of a rational agent.

### Behavioral comparison to a strategy with random choices

Despite the effect of sunk cost, it was puzzling that the fraction of optimality of 5xFR6 was, although not statistically different, slightly higher than that of 2xFR6. What strategy might be deployed to make this possible? In order to answer this question and better understand the behavior of mice, we simulated agents with different strategies performing the switching task. Among the several models and strategies that we tested, we found a plausible explanation from a strategy with random choices (see Comparison to random behavior in SI Appendix).

Depending on the total number of trials to be completed, it turns out that choosing between the PR and FR sides randomly can lead to close to optimal behavior (Fig. 4B, C, D). That is, for each cost-benefit parameter pair, acting randomly can lead to close to optimal behavior as long as the agent stops at a certain number of trials (this number is equivalent to 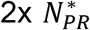, where 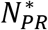 is the number of trials on the PR). For instance, the EoR of the optimality of random choices is 0.60 ± 0.01, 0.73 ± 0.02, 0.72 ± 0.02, and 0.92 ± 0.02 (mean ± SD) at 2xFR, 2xFR12, 5xFR6, and 5xFR12, respectively when the total numbers of trials were chosen from the mouse data (Fig 4C). These numbers indicate that an agent deploying a random choice strategy could collect reward efficiently enough, except in the 2xFR6 condition.

Compared to 2xFR6, the other conditions have relatively higher PR values, which encouraged mice to revisit the PR side more often and generated the resemblance of random choices. The efficiency of the mouse behavior compared to random choices was quantified (Fig. 4D). While the mice were far more efficient in 2xFR6, the other conditions were only marginally efficient compared to the agent with random choices. This analysis provided insight into the relationship between parameter values and the pattern of the behavioral outcome of mice. To summarize, the resemblance of random choices in 5xFR6 contributed partially to the high optimality despite the presence of the sunk cost fallacy.

## Discussion

Applying quantitative behavior and a theoretical framework to a mouse model system, we investigated the factors that contribute to suboptimal outcomes in an economic choice behavior. We found that both the cost and benefit differentially contributed to the suboptimality. An increased cost reduced the success rate of reward collection and was the main factor contributing to suboptimality, but the benefit parameter also contributed indirectly via sunk cost.

Our task design was inspired by several previous studies spanning the areas of foraging theory, economic decision-making, motivation, and irrationality (17, 20–26). Whether animals are rational or irrational is a controversial and active research question (28). Several previous studies with laboratory animals reported that foraging decisions often approximate the optimal solution, while others reported to the contrary. The various meanings of rationality used in different fields add additional complexity (12, 28). Because the behavioral readout of the laboratory animals is contingent on several factors such as internal state, training history, and contexts that are sometimes difficult to control, the results from one study tend to only support one side of the arguments. In this regard, a novel contribution of the current study is to provide a behavioral paradigm where both (close to) rational and irrational regimes exist in the same task.

A few things need to be discussed about the potential pitfalls in our task and interpretation of the results. First, although water and food rewards are widely used in animal studies, they are different from non-sating rewards (29, 30). Changes in the motivational level of animals due to satiation is unavoidable. Since motivation is a key factor that shapes the animals’ behavior, caution is needed when interpreting results with satiating rewards. In our task, however, the switching decisions at the indifference points occurred at the beginning of the session. Because the total number of trials that mice perform are an order of magnitude higher, we argue that the sating effect of water reward in our task is negligible. Second, the estimated indifference points in our study only approximates the true equilibrium because we did not test the hysteresis effect on the indifference point. In order to account for the hysteresis effect, the indifference points estimated from the PR to FR switching should be compared to a switching from FR to PR.. While it is very difficult to test for hysteresis in our task design, according to one study with probabilistic discounting as a cost parameter, hysteresis indeed exists in rat behavior (31).

Although our choice of the EoR (Expectation of Ratio) as an optimal model led several important conclusions in the current study (27), the limitations of this model also need to be discussed. First, according to the EoR, the total number of PR trials completed is an important factor that determines the optimal rate, but the order of choices has no effect. In our switching task, mice almost always stayed on the PR side at the beginning and switched to the FR side as the cost of PR side increased. The EoR model alone could not account for this behavioral pattern. According to our simulation results (data not shown), however, a simple reinforcement learning model where the reward is replaced with the per-trial ratio of reward to lever presses explains this behavior well. Second, revisiting the PR side even after passing the indifference point increased the number of the PR trials, which led to an increase in the fraction of optimality. Therefore, the estimated values in this study are likely to be an overestimation from or close to the upper limit of the true values. Third, effort and time cost are different physical entities and thought to be processed in different computational modules in the brain (24, 32–35). The current model neither tried to separate them, nor considered the nonlinearity of time delay and effort cost. These issues need to be addressed in future studies. Finally, other models and strategies may exist that potentially explain mouse behavior in the switching task better (3, 4, 36). Taken together, although the EoR serves as a useful reference to understanding the mouse behavior in our task, these limitations should be in mind when interpreting the results.

Based on the identification of a regime where the cost parameter and sunk cost had little impact on the suboptimality, this study supports a model whereby parallel neural circuits independently activate and modulate multiple valuation algorithms (16, 33). In the rational regime, the large reward option was particularly unattractive and the cost of the small reward option particularly low. According to this model of parallel circuits, therefore, a different circuit that processes covert factors might not be recruited into valuation in this condition. Because the systematic adjustment of the parameters can lead the transition between near-optimal and suboptimal, the application of advanced tools and techniques available in the mouse model system to our behavioral platform can make it possible to test this model directly.

Valuation is a fundamental cognitive process in decision-making and its dysfunction is expressed in diverse psychopathology including addiction, schizophrenia, depression, anxiety disorders, and severe impulsivity (37–41). Therefore, considering the parametric flexibility of this behavior, applying the approaches used in this study to disease models may serve as a useful platform to advance the understanding of biological substrates underlying rational and irrational decision factors as well as the function and dysfunction of the neural circuits involved in this process.

## Materials and Methods

### Animals

Five female and five male mice of the strain C57BL/6J were used in this study. These animals were bred on site from mice purchased from Charles River Laboratories (Wilmington, MA). The mice tested were between the ages of 8 and 13 months. All mice were kept on a 12hr/12hr light-dark cycle. All experiments were performed according to the guidelines of the Institutional Animal Care and Use Committee at Brandeis University.

### Behavioral setup and switching task

Experiments were conducted in a behavior chamber (SanWorks, Stony Brook, NY) containing nose-pokes and levers with infrared LED/infrared phototransistor pairs (Fig. 1). The chamber was connected to a Bpod state machine (SanWorks) and trial events were triggered through Matlab (MathWorks, Natick, MA). As described in the main text, the task involved combining a progressive ratio (PR) and fixed ratio (FR) schedule of lever pressing. The PR was associated with a large volume of water (either 6μL or 15μL) and the FR was associated with a small volume of water (3μL). In addition, the FR could either be 6 presses or 12 presses. At the end of each session, the mouse was weighed and additional water was given at the end of each session if necessary to maintain the animal’s weight at 85-90% of their free-drinking body weight.

### Data analysis

All data analysis was carried out using built-in and custom-built software in Matlab (Mathworks). Statistical significance among four parameter pairs was first tested with the two-factor nonparametric Scheirer-Ray-Hare test and the post-hoc pairwise comparison was done with Wilcoxon Rank-sum test with Bonferroni correction for 4 comparison pairs. For single pair comparisons, a nonparametric Wilcoxon Rank-sum test was used. A *P value* cutoff of 0.05 was used for significance testing. Sunk cost analysis was done in a similar way as described in Sweis et al (9) with the number of PR presses as a variable. To prevent data from being skewed by a single mouse, data points were only included if there were more than 5 completed trials in that condition, suggesting that at least half of the mice participated. The overall effect of sunk cost in each context was quantified using a two-way ANOVA with the probability as the dependent variable and number of presses remaining x sunk cost groups as factors. Post-hoc comparisons were done between the regression slopes of investment groups as well as each slope to an adjusted control slope (Fig. S3) using an analysis of covariance (ANOCOVA) with p-values adjusted for multiple comparisons. Calculation of EoR optimality and comparison to random behavior are described in SI Appendix.

## Acknowledgments

This work is supported by NARSAD Young Investigator Grant and NIH R01 MH110391, NIH R01 NS104818, NIH F31 DA051155. We thank the Lisman laboratory for assistance and, Drs. YS Kim and anonymous reviewers from the previous review process for comments, suggestions and ideas.

