## Supporting Information for "Parameter-dependent shift from rational to irrational decisions in mice"

**This PDF file includes:**

Supplementary text

SI References

Figures S1 to S3

Tables S1

Legends for Movies S1 to S2

**Other supplementary materials for this manuscript include the following:**

Movies S1 to S2

### Supplementary Information Text

**Maximization of different currencies yields the same optimal behavior.** In past foraging literature there have been two "currencies" that have been studies (1). The first is known as the "long-term rate" or the "ratio of expectations" (RoE) and is defined as:

$$(1) \qquad RoE = \frac{\sum_{k=1}^{N} G_k}{\sum_{k=1}^{N} T_k}$$

, where $G_k$ = the "energy gain" from the k'th food item and $T_k$ = the time spent acquiring the k'th food item. For this task, we replace the G with the size of the reward (in microliters, µl) and the T with the number of lever presses required. The index k will now index the trial number. The alternative currency is known as the "expectation of ratios" (EoR) and is defined as:

$$(2) \qquad EoR = \frac{1}{N}\sum_{k=1}^{N} \frac{G_k}{T_k}$$

Using the same variable definitions as above. Here we use these definitions to calculate the optimal number of trials the mouse should spend at each lever for a given session type (2xFR6, 2xFR12, 5xFR6, or 5xFR12).

***<u>RoE analysis.</u>*** To get the optimal behavior from an RoE perspective, we calculate, given a certain number of lever presses, what is the optimal distribution of these lever presses between the large and small reward levers where optimality means acquiring as much reward as possible. We start by writing down an equation for the reward acquired:

$$(3) \qquad R(N) = N_{FR}\left(\frac{FR}{P_{FR}}\right) + f(N_{PR}) * PR$$

, where $N = N_{FR} + N_{PR}$ (# of presses at fixed + # of presses at progressive reward side), FR = fixed reward size, PR = progressive reward size, $P_s$= # of presses required at the FR lever for reward, and $f(N_{PR})$ = # of trials corresponding to $N_{PR}$ lever presses at the PR lever. Note that we assume the optimal strategy does not contain any aborted trials since an aborted trial will always push RoE towards 0. To start, we need to figure out what $f(N_{PR})$ is. To do this, we set up the following equality

$$N_{PR} = \sum_{k=1}^{f(N_{PR})} k + 1$$

Expressing that the number of PR presses increases by one with each additional successful PR trial. We then evaluate the sum and solve for $f(N_{PR})$.

$$0 = f(N_{PR})^2 + 3f(N_{PR}) - 2N_{PR}$$

(4) $$f(N_{PR}) = \frac{1}{2}\left(\sqrt{8 * N_{PR} + 9} - 3\right)$$

Here, we ignore the other root since it is strictly negative. We now plug Eq 4 into Eq 3 and replace $N_{FR}$ with $N - N_{PR}$:

$$R(N) = (N - N_{PR})\left(\frac{FR}{P_{FR}}\right) + \frac{1}{2}(\sqrt{8 * N_{PR} + 9} - 3) * PR$$

Now we want to take the derivative of $R(N)$ w.r.t $N_{PR}$ and solve for the $N^*_{PR}$ that will give us $R^*(N)$ (the maximum reward achievable given N lever presses).

$$\frac{dR(N)}{dN_{PR}} = \frac{2PR}{\sqrt{8 * N_{PR} + 9}} - \frac{FR}{P_{FR}}$$

Solving for $N_{PR}$ gives:

(5) $$N^*_{PR} = \frac{1}{8}\left(\left(\frac{2PR * P_{FR}}{FR}\right)^2 - 9\right)$$

Note that Eq 5 yields an $N^*_{PR}$ that is independent of N.

For the 2xFR6 case (FR = 3, PR = 6, $P_{FR}$ = 6) this yields $N^*_{PR}$= 70.875 presses, $f(N^*_{PR})$ = 10.5 trials. For the 2xFR12 case (FR = 3, PR = 6, $P_{FR}$ = 12) this yields $N^*_{PR}$ = 286.875 presses, $f(N^*_{PR})$ = 22.5 trials. For the 5xFR6 case (FR = 3, PR = 15, $P_{FR}$ = 6) this yields $N^*_{PR}$ = 448.875 presses, $f(N^*_{PR})$ = 28.5 trials. For the 5xFR12 case (FR = 3, PR = 15, $P_{FR}$ = 12) this yields $N^*_{PR}$ = 1798.875 presses, $f(N^*_{PR})$ = 58.5 trials.

***EoR analysis.*** The EoR for a given number of trials $N$ is:

(6) $$EoR(N) = \frac{1}{N}\sum_{k=1}^{N}\frac{r_k}{p_k}$$

Where $N$ is the total number of trials, $r_k$ is the reward (in $\mu l$) received on trial k, and $p_k$ is the number of lever presses done on trial k. Note that this is different than the typical EoR discussed in past studies since the denominator of each fraction is in terms of an energetic cost instead of a time cost. We are now interested in calculating, for each session type (2xFR6, 2xFR12, 5xFR6, 5xFR12) what the optimal number of trials at the progressive reward is ($N_{PR}^{*}$). To do this we want to find which $N_{PR}$ gives us the highest $EoR$ for a given $N$. We can find this by splitting the above equation for $EoR$ up into terms for trials at the PR and trials at the FR.

(7) $$EoR(N) = \frac{1}{N}\left(r_{PR} * \sum_{k=1}^{N_{PR}}\frac{1}{k+1} + N_{FR} * \frac{r_{FR}}{p_{FR}}\right)$$

Where $r_{PR}$ is the reward from the PR side, $r_{FR}$ is the reward from the FR side, $p_{FR}$ is the number of lever presses required for reward at the FR, and $N_{FR}$ is the number of completed trials at the FR side. Note that we assume that an optimal agent does not abort any trials since this would strictly decrease the $EoR$ towards 0. Substituting $N_{FR} = N - N_{PR}$ and evaluating the sum we get

(8) $$EoR(N) = \frac{1}{N}\left(r_{PR} * \left(\psi^{(0)}(N_{PR}+2) + \gamma - 1\right) + (N - N_{PR}) * \frac{r_{FR}}{p_{FR}}\right)$$

Where $\psi^{(0)}(x)$ is the digamma function and $\gamma$ is the Euler-Mascheroni constant. To find $N_{PR}^{*}$ we take the derivative of this expression w.r.t $N_{PR}$

(9) $$\frac{dEoR(N)}{dN_{PR}} = \frac{1}{N}\left(r_{PR} * \psi^{(1)}(N_{PR}+2) - \frac{r_{FR}}{p_{FR}}\right)$$

Where $\psi^{(1)}(x)$ is now the trigamma function (derivative of digamma). Setting the derivative to 0 and solving for $\psi^{(1)}(N_{PR}+2)$ we get

(10) $$\psi^{(1)}(N_{PR}^{*}+2) = \frac{r_{FR}}{r_{PR} * p_{FR}}$$

from which we can get numerical approximations for $N_{PR}^{*}$ which are 10.5, 22.5, 28.5, and 58.5 at 2xFR6, 2xFR12, 5xFR6, and 5xFR12, respectively. These numbers have a straightforward interpretation. For example, on the 10'th PR trial in a 2xFR6 session, the mouse will receive 6$\mu l$ of water for 11 lever presses which still gives a better ratio than the FR side ($3\mu l$ for 6 presses). On the 11'th PR trial these ratios will be equal (6/12 and 3/6). This pattern is true for all 4 session types. Therefore the .5 on each of the above $N_{PR}^{*}$ reflects that fact that it is equivalent to stop going to the PR side either when the ratios become equal or 1 trial before. For the analysis in this paper, we took the optimal $N_{PR}^{*}$ for each session type respectively to be 10, 22, 28, and 58. To then calculate each mouse's fraction of EoR optimality for each session, we calculated the mouse's observed EoR for that session (mean of per-trial reward divided by the number of presses, aborted trials included) and divided that by the optimal EoR given the number of total trials the mouse performed (including aborted trials) and the session type.

Taken together, analysis of both potential currencies give the same answer (to within numerical error).

**Comparison to random behavior.** Another interesting perspective from which to analyze the mice's behavior is from the perspective of a randomly choosing agent. This tells us to what degree the mice's choices reflect decisions beyond simple random choice between sides. Interestingly, random choice can lead to optimal or close to optimal behavior for a range of total trials completed. That is, if a randomly choosing agent performs 2x$N_{PR}^{*}$ trials, then on average it will choose the PR side $N_{PR}^{*}$ times, resulting in an optimal EoR. To compute optimality distributions produced by random behavior for each session type (Figures 4A,C), we simulated 10,000 randomly choosing agents for each number of total trials from 1 to 1000. On each simulated trial, there was a 50/50 chance of choosing the FR or PR side. The observed EoR of random choices for a given number of trials N was then compared to the optimal EoR for that number of trials to get a fraction of EoR optimality of random behavior. To then compare the mice's performance to random, for each session we looked at the ratio of the observed EoR optimality of the mouse to the mean of the EoR optimality distribution of the random agents corresponding to the same number of trials the mouse had completed (Figure 4D). These ratios give how much better a mouse's choices were for a particular session type and number of trials, compared to random choice.

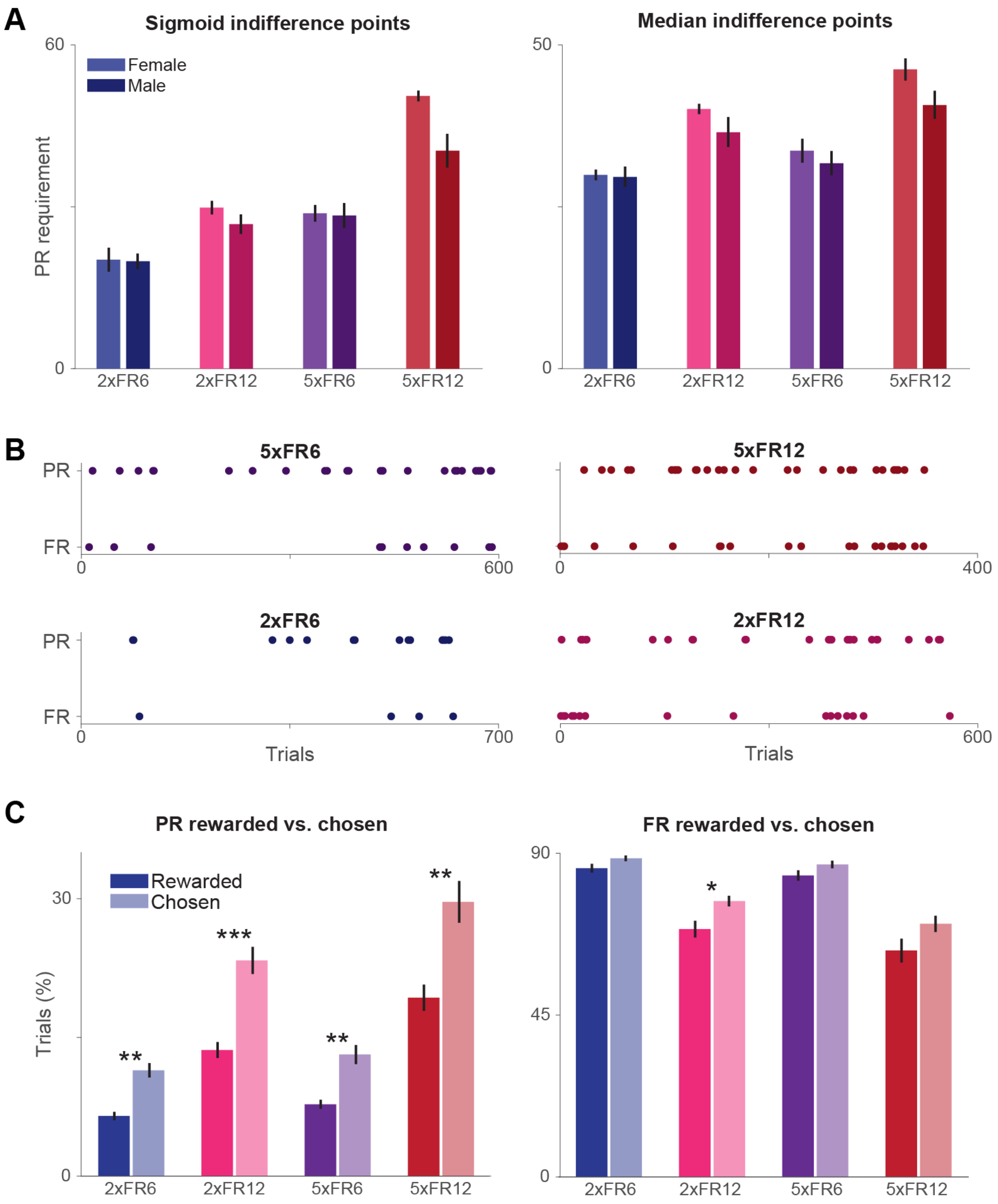


**Fig. S1. Comparison of choice behavior in various contexts. A.** Male and female mice perform similarly on the optimal switching task. When comparing indifference points of male (n = 5) and female (n = 5) mice, no significant differences were identified. As a result, male and female mice were grouped together for all other analyses. Error bars reflect standard error of the mean. **B.** Incomplete trials occur more frequently in higher value sessions. Shows example sessions for each parameter pair. Each point is a single incomplete trial within the session. All sessions are from the same animal. **C.** Mice are more likely to attempt and fail when pressing PR. Incomplete trials occur when attempting both the PR and FR. However, there is a significantly larger difference between the percentage of trials attempted on the PR and those actually rewarded (*$p < 0.05$, **$p < 0.01$, ***$p < 0.001$, Wilcoxon Rank-sum test, n = 10 mice). Error bars reflect standard error of the mean.

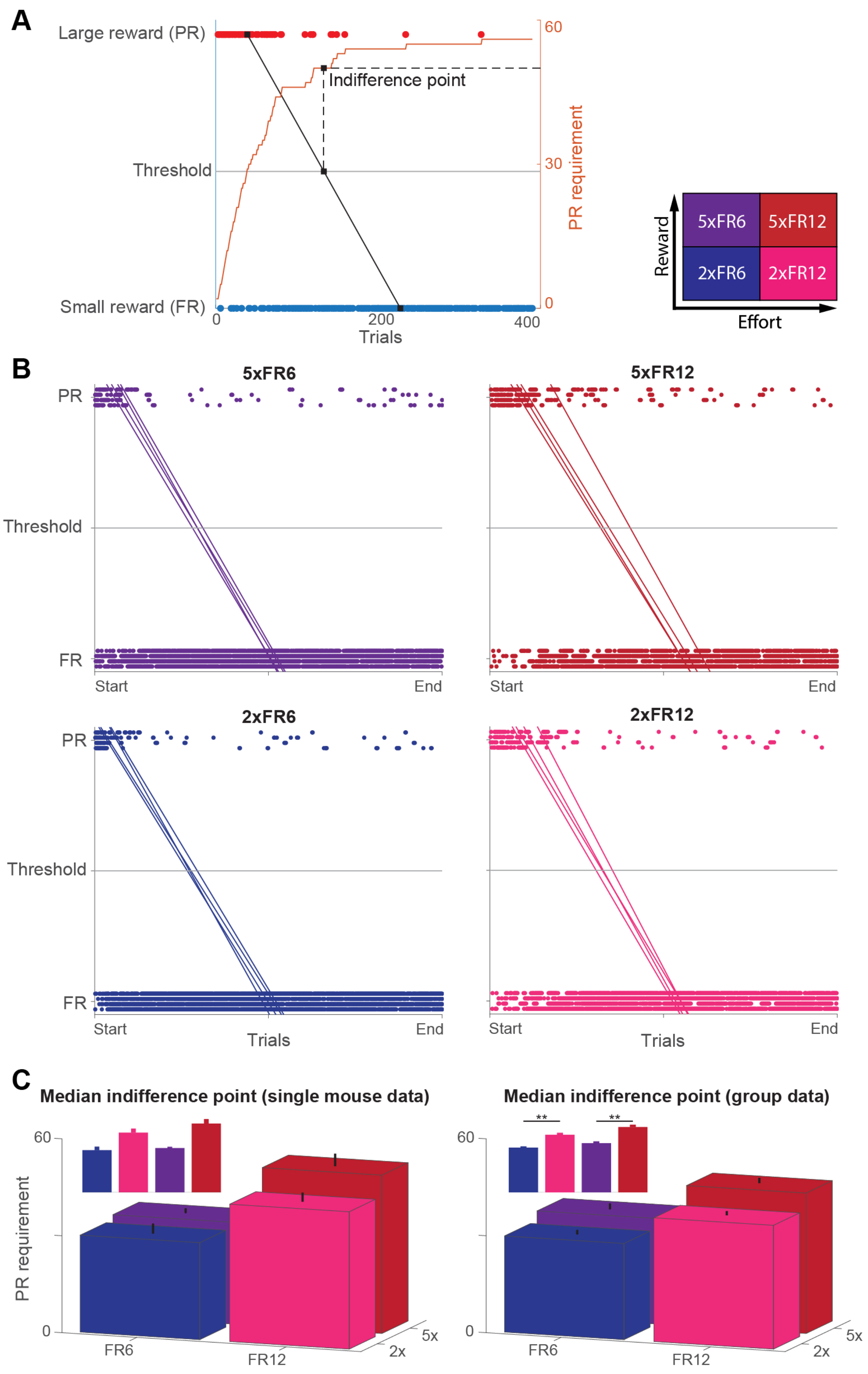
A
Large reward (PR)
Threshold
Small reward (FR)
0
200
400
Trials
Indifference point
60
30
0
PR requirement
Reward
Effort
5xFR6
5xFR12
2xFR6
2xFR12
B
5xFR6
5xFR12
2xFR6
2xFR12
PR
Threshold
FR
Start
End
Trials
C
Median indifference point (single mouse data)
Median indifference point (group data)
PR requirement
60
0
**
FR6
FR12
5x
2x

**Fig. S2. Calculating indifference points using median trial values displays variation in switching behavior between parameter conditions. A**. Example session showing how the indifference point is estimated. Black line is the linear fit and orange curve is the cumulative PR requirement over time. First, the median trial numbers for PR and FR trials are calculated and connected by a line. Then, where the line crosses the threshold, a trial number is identified (black squares). Finally, the indifference point is determined by finding the number of presses required for the PR at this trial (dashed lines). **B**. Median lines for all 16 sessions of one mouse, sorted by session parameters. Trials are plotted as individual points normalized to the total number of trials in the session and each row of points is a different session. **C**. Indifference points for a single mouse (left, n = 4 sessions) and the entire population (right, n = 10 mice). Error bars reflect standard error of the mean. A two-way Scheirer-Ray-Hare test indicated a significant effect in the FR requirement ($H_1 = 22.70$, $p = 2 \times 10^{-6}$). Insert shows the same data with significant pairs notated (post hoc Wilcoxon Rank-sum test with Bonferroni correction, $^{**}p < 0.01$).

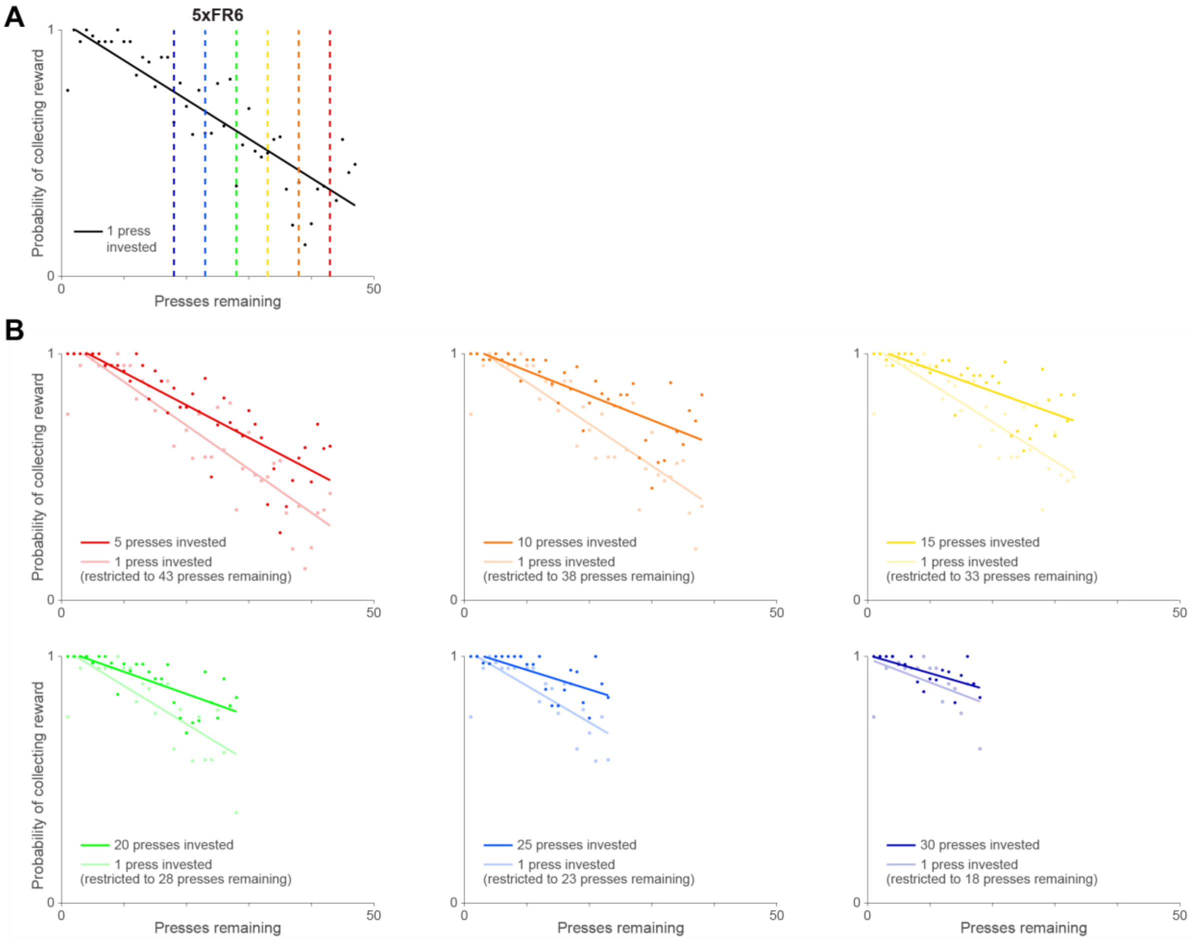


**Fig. S3. Visual representation of adjusted control comparisons in sunk cost analysis.** In order to correct for variations in the amount of data available for each sunk cost group, we calculated an "adjusted control" to which we could compare each group. This consisted of taking the data points of the 1 press invested group and using only those data points whose range overlapped with the sunk cost group being compared. For example, in our 5xFR6 context, the 1 press invested group has data points that extend up to 47 presses remaining; however, the 5 presses invested group only extends to 43 presses remaining. Therefore, we take the regression of the full dataset for the 5 presses invested and compare it to a new regression of the 1 press invested group restrained to 1-43 presses remaining. **A.** Control condition (1 press invested) data points and linear fit in the 5xFR6 context. Dashed colored lines represent the highest press remaining that each sunk cost group extends. Thus, only the data points between the y-axis and the corresponding dashed line will be used in the adjusted control. **B.** Data points and regressions for each sunk cost group plotted against a new adjusted control regression calculated based on the 1 press invested points limited to the overlapping data range.

**Table S1. Trials and rewards for each parameter.**

| | 2xFR6 | 2xFR12 | 5xFR6 | 5xFR12 |
|---|---|---|---|---|
| **Total Trials** | 590 ± 120 | 380 ± 60 | 540 ± 80 | 320 ± 100 |
| **Completed Trials** | 550 ± 130 | 310 ± 60 | 500 ± 90 | 270 ± 100 |
| **Reward Collected (μL)** | 1700 ± 400 | 1100 ± 200 | 1900 ± 300 | 1400 ± 300 |

Shows the average and standard deviation (n = 10 mice) of total number of trials, number of trials where reward was collected, and the total amount of reward collected. Fewer trials were done and less reward was collected in the FR12 conditions, which is likely due to the increased amount of time it takes to press the lever 12 times.

**Legends for Movies**

**Movie S1.** An example of a mouse pressing a lever on the FR side with a small reward. The mouse must press 6 times to collect reward.

**Movie S2.** An example of a mouse pressing a lever on the PR side with a large reward. Three consecutive trials were presented. The mouse must first press 2, then 3, then 4 times to collect reward.
